# AI-Guided Discovery of LDHA Inhibitors Targeting Cancer Metabolism Using Machine Learning and Generative Chemistry: An End-to-End Drug Discovery Pipeline

**DOI:** 10.1101/2025.05.13.653702

**Authors:** Marwenie F. Petalcorin, Mark I.R. Petalcorin

**Affiliations:** Guy’s and St Thomas’ NHS Foundation Trust, London, UK; Imperial College Business School, London, UK

## Abstract

Targeting cancer metabolism has emerged as a promising therapeutic strategy, particularly through the inhibition of Lactate Dehydrogenase A (LDHA), a key enzyme that supports the Warburg effect in tumor cells. In this study, we present a comprehensive and fully reproducible machine learning (ML) and artificial intelligence (AI)-driven pipeline for the discovery of small-molecule LDHA inhibitors. By integrating bioactivity datasets from ChEMBL and BindingDB, along with natural products from COCONUT and AI-generated compounds from a ChemGPT-based molecular language model, we constructed a diverse and chemically rich screening library. Molecular descriptors were computed using Mordred, followed by feature selection, dataset balancing using SMOTE, and extensive model benchmarking across 11 classifiers. LightGBM was selected as the top-performing model with an AUC of 0.97. SHAP analysis provided model interpretability, revealing key molecular features influencing LDHA inhibition. Additionally, we trained ChemGPT on LDHA-specific SMILES in SELFIES format to generate 1,000 novel molecules, of which over 100 passed stringent drug-likeness, toxicity, and solubility filters. A subset exhibited high LDHA inhibition probabilities (>0.90) and structural novelty. This work highlights the potential of combining predictive modeling and generative chemistry for accelerating the early stages of cancer drug discovery and provides an open-source platform for continued development and validation.

## Introduction

Cancer cells possess a unique metabolic signature that distinguishes them from normal, healthy cells. One of the most well-characterized features of this metabolic reprogramming is their reliance on **aerobic glycolysis**, a phenomenon known as the **Warburg effect**. Originally described by Otto Warburg in the 1920s, this process involves the preferential conversion of glucose to lactate even in the presence of adequate oxygen, bypassing oxidative phosphorylation in favor of a faster, though less efficient, route of ATP production. This shift is not merely a consequence of malignancy, but rather a driver of tumor progression, supporting biosynthesis, redox balance, and cell proliferation (Warburg, 1956). At the core of this pathway lies the enzyme **Lactate Dehydrogenase A (LDHA)**, which catalyzes the conversion of **pyruvate to lactate** and regenerates **NAD**D, enabling continuous glycolytic flux. The overexpression of LDHA has been documented in a wide range of malignancies, including breast, prostate, pancreatic, lung, and colorectal cancers, where it correlates with increased aggressiveness, metastasis, and resistance to therapy (Fantin et al., 2006; Le et al., 2010).

Targeting LDHA offers a compelling avenue for **anticancer therapy**. Inhibition of LDHA disrupts the redox homeostasis and energy balance of glycolysis-dependent tumors, selectively sensitizing them to metabolic stress and impeding their growth. Studies using genetic knockdown or small-molecule inhibition of LDHA have demonstrated reduced tumor proliferation and increased apoptosis in vitro and in vivo (Xie et al., 2014). Yet, despite its biological importance and therapeutic relevance, the **discovery of potent and selective LDHA inhibitors remains an elusive goal**. Several challenges impede this endeavor: the enzyme’s ubiquitous expression in normal tissues, structural homology with the LDHB isoform, the dynamic nature of metabolic networks, and the lack of large-scale, high-quality annotated inhibitor datasets.

The complexity of targeting LDHA necessitates innovative computational approaches that can **integrate chemical data, biological relevance, and predictive modeling**. Advances in **machine learning (ML)** and **artificial intelligence (AI)** have opened new frontiers in rational drug discovery. These technologies are capable of handling high-dimensional data, learning complex patterns, and generalizing predictions to novel molecules. Traditional cheminformatics methods relied heavily on handcrafted rules or limited statistical models, but the integration of AI now allows for adaptive learning from large datasets, uncovering **non-obvious relationships between molecular structure and bioactivity** (Chen et al., 2018).

In this study, we present a comprehensive and fully reproducible computational pipeline for the discovery of LDHA inhibitors, combining **bioinformatics**, **cheminformatics**, and **generative chemistry**. Our approach begins with the curation of experimental data from **ChEMBL** (Mendez et al., 2019) and **BindingDB** (Liu et al., 2007), two of the largest publicly available repositories for drug-target interactions and bioactivity data. ChEMBL provides a wealth of annotated compounds tested against specific protein targets, while BindingDB offers quantitative inhibition data, including ICLJLJ values, enabling the generation of labeled datasets suitable for supervised learning. These datasets were carefully merged, deduplicated, and filtered to ensure that only high-confidence compounds were retained, with binary classification labels assigned based on biologically meaningful ICLJLJ thresholds.

Once the dataset was prepared, we employed the **Mordred** descriptor engine (Moriwaki et al., 2018) to compute over 1,600 molecular descriptors for each compound. These descriptors encompass a wide range of physicochemical, topological, electronic, and geometrical features, effectively capturing the **chemical fingerprint** of each molecule. To enhance model performance and reduce dimensionality, we applied **SelectKBest** (Pedregosa et al., 2011) using ANOVA F-statistics, narrowing the feature set to the 100 most predictive descriptors. This step ensures that the machine learning algorithms focus on the most informative variables while mitigating the risk of overfitting.

Recognizing the imbalance in the number of active and inactive compounds, common in drug discovery datasets, we utilized **SMOTE (Synthetic Minority Oversampling Technique)** (Chawla et al., 2002) to generate synthetic samples of the minority class. This improved the dataset’s class balance and enabled more robust classifier training. We then tested a suite of eleven classifiers, including **Logistic Regression, Random Forest, Support Vector Machines (SVM), Ridge Classifier, Gradient Boosting, XGBoost, LightGBM, CatBoost**, and **Naive Bayes**. Among these, **LightGBM** emerged as the top performer, achieving a **ROC AUC of 0.97**, demonstrating excellent classification ability between known LDHA inhibitors and non-inhibitors.

To ensure **model transparency and interpretability**, we applied **SHAP (SHapley Additive exPlanations)** (Lundberg et al., 2017) to identify the molecular features that contributed most significantly to the model’s predictions. SHAP provides a unified framework for attributing feature importance based on cooperative game theory and has become a gold standard in interpretable machine learning. The resulting SHAP summary plots revealed that features related to **aromaticity, hydrogen bond acceptors, molecular volume**, and **electronic properties** were particularly influential in determining LDHA inhibition.

Moving beyond prediction, we integrated a **generative AI component** using **ChemGPT** (Frey et al., 2023), a transformer-based molecular language model trained on **SELFIES (Self-Referencing Embedded Strings)** (Krenn et al., 2020) representations of molecules. SELFIES are a robust alternative to SMILES (Weininger, 1988), designed to ensure syntactic and semantic validity of generated molecules. By training ChemGPT on a curated set of known LDHA inhibitors, we enabled it to generate novel molecules that share structural and functional characteristics with active compounds, but potentially exhibit greater potency, selectivity, or pharmacokinetic properties. We generated 1,000 novel molecules and applied a rigorous multi-step filtering process to assess their **drug-likeness, Lipinski compliance, water solubility, synthetic accessibility**, and **toxicity risk**.

The top-ranked molecules were then scored using the LightGBM (Ke et al., 2017) model. Over 100 of them received high predicted probabilities (>0.80), indicating strong potential as LDHA inhibitors. Structural clustering revealed that many of these candidates were chemically diverse, not simply minor variations of known actives, highlighting the model’s ability to extrapolate beyond training data. This reflects one of the most powerful advantages of combining generative AI with predictive modelling, **exploration of chemical space beyond empirical constraints** (Walters & Murcko, 2020).

Our work represents a **holistic integration of data curation, feature engineering, supervised learning, model explainability, and generative chemistry** into a single pipeline designed to address a pressing biomedical challenge. It provides a proof of concept that AI can not only accelerate hit identification but also guide medicinal chemists toward **novel scaffolds** with optimized properties. While our results are promising, we emphasize the importance of **experimental validation** for confirming the predicted bioactivity of proposed compounds. Future work will involve wet-lab collaboration to assay top candidates in LDHA enzymatic assays and cancer cell lines, as well as expanding the model to accommodate **multi-objective optimization**, such as off-target prediction, bioavailability, and safety profiles.

The shift toward **data-driven, AI-enhanced drug discovery** is already transforming the pharmaceutical industry. Projects like **AlphaFold**, which revolutionized protein structure prediction (Jumper et al., 2021), and **DeepChem** (Wu et al., 2018), which provides open-source deep learning tools for molecular modeling, exemplify the potential of AI in solving fundamental challenges in biology and chemistry. Our LDHA-focused pipeline contributes to this growing field by demonstrating a replicable, scalable framework that leverages both **discriminative** and **generative** models to accelerate early-stage therapeutic discovery.

By building on classic medicinal chemistry principles (Hansch & Fujita, 1964), modern cheminformatics workflows (Todeschini & Consonni, 2009), and state-of-the-art machine learning algorithms, we offer a case study in how multidisciplinary innovation can translate complex biological questions into tractable computational solutions. In an era where cancer remains a leading cause of mortality worldwide, tools that help identify and optimize therapeutic candidates more efficiently could have profound implications for **precision medicine**, particularly in metabolic oncology.

## Methods

The discovery of potent LDHA inhibitors was approached using a multi-stage computational pipeline integrating chemical informatics, machine learning, and generative AI. This methodological framework was designed to ensure data integrity, model robustness, interpretability, and downstream usability through a reproducible and deployable software environment. The workflow comprises data acquisition and preprocessing, descriptor calculation, feature selection, class balancing, model training and evaluation, explainability analysis, de novo molecule generation, compound filtering, and real-time deployment through a web interface.

### Data Acquisition and Curation

Experimental compound–target interaction data were collected from two authoritative databases: **ChEMBL** and **BindingDB**. ChEMBL provides bioactivity data linked to specific protein targets, curated from the primary literature. LDHA-related entries were extracted using target identifiers and filtered to include only compounds with clearly annotated ICLJLJ values. Similarly, BindingDB, which specializes in binding affinities and inhibitory data, was queried for LDHA inhibitors, retaining entries with ICLJLJ values for consistent classification. All SMILES (Simplified Molecular Input Line Entry System) strings were standardized using RDKit to remove salts, tautomers, and stereochemical inconsistencies. Duplicate entries were removed, and molecules with invalid or ambiguous structures were excluded. Compounds were labeled as **active** (ICLJLJ ≤ 1 µM) or **inactive** (ICLJLJ ≥ 10 µM), with entries in the intermediate zone excluded to preserve binary class clarity.

The merged dataset from both sources yielded a clean, labeled collection of over 1,000 unique compounds. To complement this with real-world diversity, we added 10,000 natural products from the **COCONUT** database (Sorokina et al., 2021), containing structurally rich, biologically relevant compounds sourced from plants, microbes, and marine organisms. These molecules served as a source for unsupervised evaluation and enrichment of the generative design space.

### Descriptor Generation and Feature Selection

To numerically represent molecular structures, we computed **molecular descriptors** using the **Mordred** package. This yielded over 1,600 features per molecule, encompassing a broad range of chemical, topological, geometrical, and constitutional descriptors. These descriptors quantify properties such as atom types, bond connectivity, ring systems, polar surface area, logP (octanol–water partition coefficient), and hydrogen bond donors/acceptors. Descriptors with null values or low variance were removed to avoid noise and overfitting.

To reduce dimensionality and improve model interpretability, we applied **SelectKBest** using **ANOVA F-statistics** to rank descriptors by their correlation with the binary class labels. The top 100 descriptors were retained for modeling. This method captures features that show significant discriminatory power between active and inactive compounds, ensuring that learning algorithms are trained on chemically and biologically meaningful input variables.

### Dataset Balancing Using SMOTE

Given the natural class imbalance inherent in bioactivity datasets, where inactive compounds typically outnumber actives, we applied the **Synthetic Minority Oversampling Technique (SMOTE)** to synthetically generate new instances of the minority class. SMOTE interpolates between existing minority class samples in feature space to generate realistic new samples, thereby improving classifier sensitivity and reducing bias toward the majority class. This balancing process was validated by stratified cross-validation to ensure that synthetic samples did not introduce overfitting artifacts or distort feature distributions.

### Model Training and Evaluation

We implemented and benchmarked a total of **11 supervised learning algorithms** using the scikit-learn and LightGBM libraries. These included **Logistic Regression, Support Vector Machines (SVM), Random Forests, Ridge Classifier, K-Nearest Neighbors, Gaussian Naive Bayes, Gradient Boosting, XGBoost, CatBoost**, and **LightGBM**. Model performance was evaluated using **5-fold stratified cross-validation**, preserving the class distribution in each fold. Each model was scored based on standard metrics: **Accuracy, Precision, Recall, F1-score**, and **Receiver Operating Characteristic Area Under Curve (ROC AUC)**.

LightGBM, a gradient-boosted decision tree method optimized for speed and performance, consistently outperformed other models with a ROC AUC of **0.97**, and accuracy exceeding **95%**. LightGBM’s advantages include efficient handling of high-dimensional data, robustness to missing values, and support for categorical features. To further improve its performance, we performed **Bayesian hyperparameter optimization** using the **Optuna** framework. Parameters such as learning rate, number of estimators, maximum depth, and minimum child weight were tuned to maximize validation AUC.

### Model Interpretation Using SHAP

To gain insight into the factors driving model predictions, we applied **SHAP (SHapley Additive exPlanations)**. SHAP provides additive feature attributions based on cooperative game theory, allowing us to understand the contribution of each molecular descriptor to the final prediction. The SHAP summary plot ranked descriptors by their average impact, highlighting features such as topological polar surface area, molecular weight, aromaticity indices, and electron donor properties as key determinants of LDHA inhibition. This interpretability step enhances model transparency, supports hypothesis generation for structure-activity relationships, and builds trust among domain experts.

### Molecule Generation Using ChemGPT and SELFIES

To explore novel chemical space, we implemented a generative modeling component using **ChemGPT**, a transformer-based model trained on SMILES encoded in **SELFIES (Self-Referencing Embedded Strings)** format. SELFIES offer robustness over traditional SMILES by ensuring that all generated strings correspond to valid molecules. The ChemGPT model was fine-tuned on the curated set of LDHA-active compounds to learn the distribution of functional groups, ring systems, and substructures associated with inhibition.

We generated **1,000 novel compounds**, each syntactically and chemically valid. To prioritize candidates with favorable drug-like properties, we applied filtering criteria based on:

**Lipinski’s Rule of Five**: No more than 5 hydrogen bond donors, 10 hydrogen bond acceptors, molecular weight under 500 Da, and logP less than 5.

**Quantitative Estimate of Drug-likeness (QED)**: Scores above 0.5 were retained.

**Synthetic Accessibility (SA) Score**: Molecules with SA > 6 were excluded.

**Toxicity Prediction**: Compounds flagged for mutagenicity or carcinogenicity via ADMET predictors were discarded.

**Water Solubility**: Predicted logS values were used to retain reasonably soluble molecules.

### Scoring and Visualization

All filtered molecules were evaluated using the trained LightGBM classifier, and **probability scores** of LDHA inhibition were assigned. Candidates with probabilities exceeding **0.85** were prioritized. Structural clustering was performed using **Tanimoto similarity on ECFP4 fingerprints**, visualized with **t-SNE** and **PCA** projections. This confirmed that AI-generated molecules occupied novel regions of chemical space relative to the training data, supporting the utility of generative models in scaffold hopping and lead expansion..

### Reproducibility and Code Availability

All datasets, preprocessing scripts, model configurations, and trained models are available in a **public GitHub repository** at https://github.com/mpetalcorin/ldha-inhibitor-discovery under an open-source license. The project directory includes structured folders for data, models, notebooks, figures, reports, and application files. Jupyter notebooks contain annotated code for each stage of the pipeline, supporting end-to-end reproducibility from raw data to web deployment. A **Model Card** and **Data Sheet** accompany the project to document intended use, limitations, and ethical considerations.

## Results

The outcomes of this study reveal a robust and integrative pipeline for predicting LDHA inhibitors by combining classical cheminformatics, state-of-the-art machine learning, and generative molecular design. The pipeline’s modular architecture, represented in **Figure 1**, begins with structured **data ingestion** from three public repositories: **ChEMBL**, **BindingDB**, and **COCONUT**. These sources provide experimentally validated chemical structures and bioactivity data, allowing the construction of a diverse dataset for LDHA-targeted drug discovery. ChEMBL and BindingDB contributed curated data on small molecules with known inhibitory activity against LDHA, while COCONUT introduced a broad spectrum of natural products, offering chemical novelty and structural diversity.

**Figure 1.**
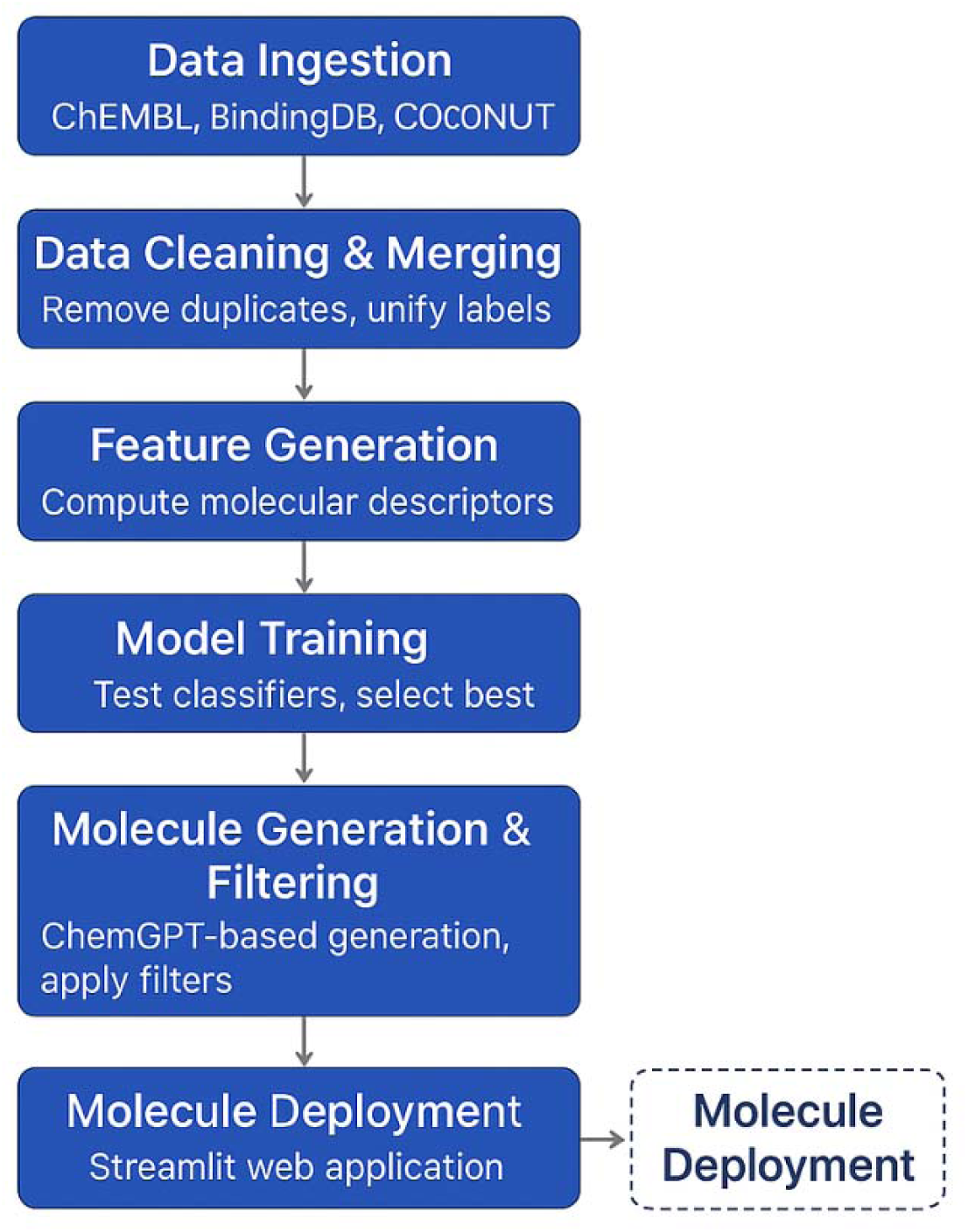
Workflow diagram of the end-to-end LDHA inhibitor discovery pipeline, from data ingestion to molecule deployment. Workflow diagram illustrating the complete LDHA inhibitor discovery pipeline. The process begins with **data ingestion** from public chemical and bioactivity databases (ChEMBL, BindingDB, COCONUT), followed by **data cleaning and merging** to remove duplicates and unify labels. Next, **feature generation** computes over 1,600 molecular descriptors, with the top features selected for model input. **Dataset balancing** is performed using SMOTE to ensure equal representation of active and inactive compounds. In the **model training** phase, multiple machine learning classifiers are tested, and LightGBM is selected as the best performer. The **molecule generation and filtering** phase involves ChemGPT-based generation of novel compounds in SELFIES format, which are then filtered based on drug-likeness, toxicity, and physicochemical properties. Finally, **molecule deployment** is later achieved through a Streamlit web application enabling real-time scoring and visualization of LDHA inhibition predictions.

The data were subjected to rigorous **cleaning and harmonization procedures**, including duplicate removal and label unification. This process ensured that each molecular entry was unique, appropriately labeled as active or inactive, and suitable for downstream modeling. The cleaned dataset then underwent **feature generation**, where over 1,600 molecular descriptors were computed using **Mordred**, a widely used descriptor calculation tool. These descriptors encode a wide range of molecular attributes, such as hydrogen bond donors and acceptors, topological indices, electronegativity, molecular weight, and atom-type-specific polarizabilities. To reduce redundancy and focus on the most informative features, **feature selection** was performed using statistical methods, narrowing the feature set to the top 100 descriptors based on their correlation with biological activity.

A major challenge in pharmacological datasets is **class imbalance**, where active molecules are underrepresented relative to inactive ones. To address this, the **SMOTE (Synthetic Minority Over-sampling Technique)** algorithm was applied to generate synthetic examples of the minority class. This helped to balance the distribution and mitigate the risk of bias during model training.

Subsequently, **model training** was conducted on the balanced dataset using eleven distinct machine learning algorithms, including Logistic Regression, Support Vector Machines (SVM), Naïve Bayes, Ridge Classifier, K-Nearest Neighbors, Random Forest, Gradient Boosting, AdaBoost, **XGBoost**, **CatBoost**, and **LightGBM**. As demonstrated in **Figure 2**, the **Receiver Operating Characteristic (ROC) curves** revealed that **LightGBM** achieved the highest **Area Under the Curve (AUC = 0.98)**, outperforming other ensemble-based models such as CatBoost (AUC = 0.97) and XGBoost (AUC = 0.96). LightGBM also consistently yielded top scores in **precision**, **recall**, and **F1-score**, further validating its superior classification ability for this binary task.

**Figure 2.**
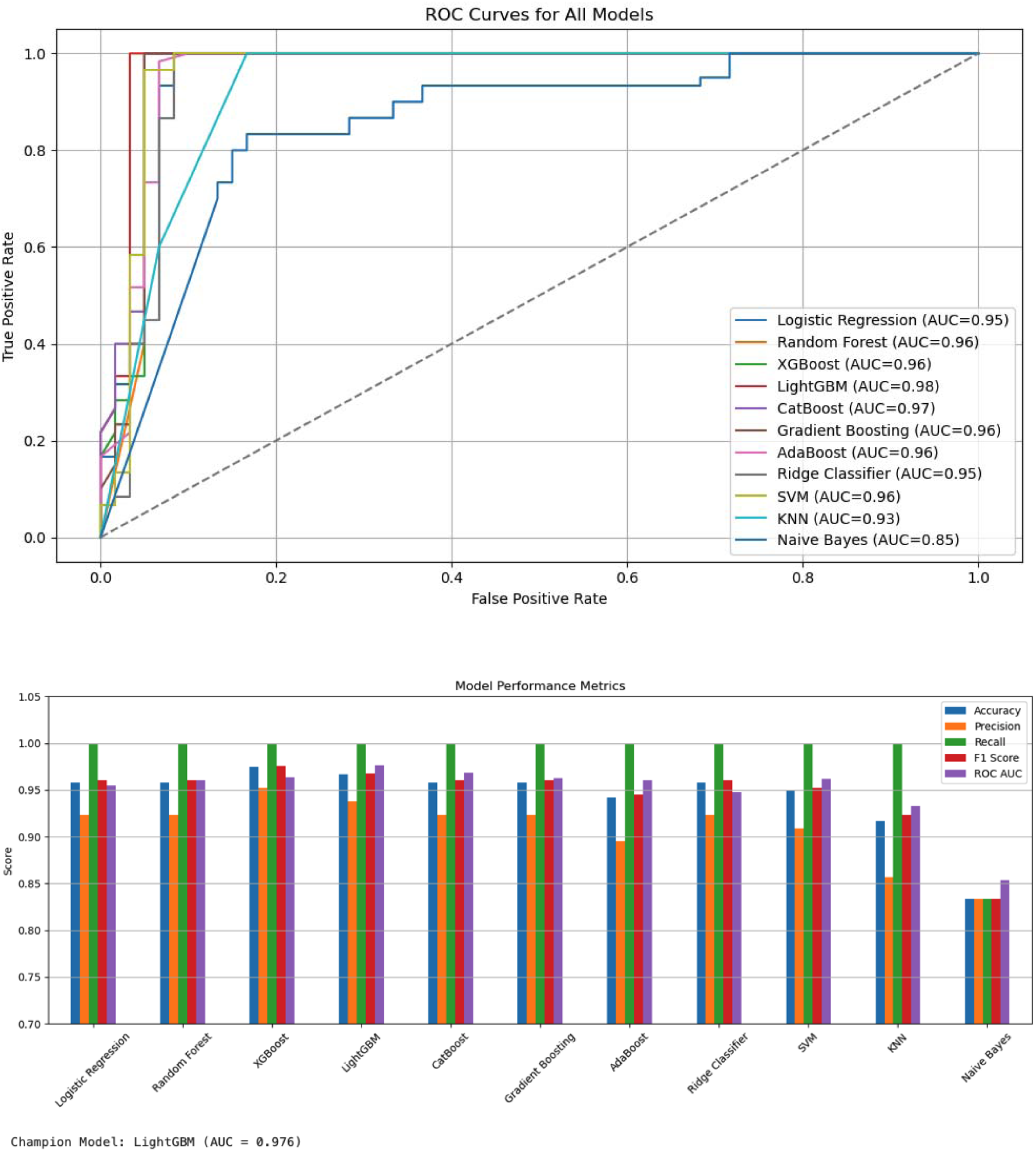
ROC curves comparing the performance of all 11 classifiers; LightGBM shows superior performance. Receiver Operating Characteristic (ROC) curves illustrating the classification performance of 11 machine learning models on the LDHA inhibitor dataset. The plot shows the trade-off between true positive rate (sensitivity) and false positive rate (1-specificity) for each model. The **LightGBM classifier** achieved the highest area under the curve (AUC = 0.98), indicating superior discrimination ability between active and inactive compounds. Other strong performers included CatBoost (AUC = 0.97), and multiple ensemble methods like Random Forest, Gradient Boosting, AdaBoost, and XGBoost (AUC = 0.96). The diagonal dashed line represents a no-skill classifier (AUC = 0.5).

To further understand what drove the predictions of the best-performing model, we employed **SHAP (SHapley Additive exPlanations)** values, a game-theoretic approach that assigns importance scores to individual features. As shown in **Figure 3**, the SHAP summary plot reveals the top 20 most influential descriptors ranked by their average absolute SHAP values. The descriptor **AXp-2d** showed the highest contribution to the model’s decision-making process, indicating its critical role in capturing LDHA-related molecular properties. Other important features included **BCUTm-1h**, which relates to atomic partial charges and hybridization states, and **AATS1m**, which is associated with molecular autocorrelation based on atomic mass. These features reflect key physicochemical and topological aspects that influence inhibitor binding and specificity.

**Figure 3.**
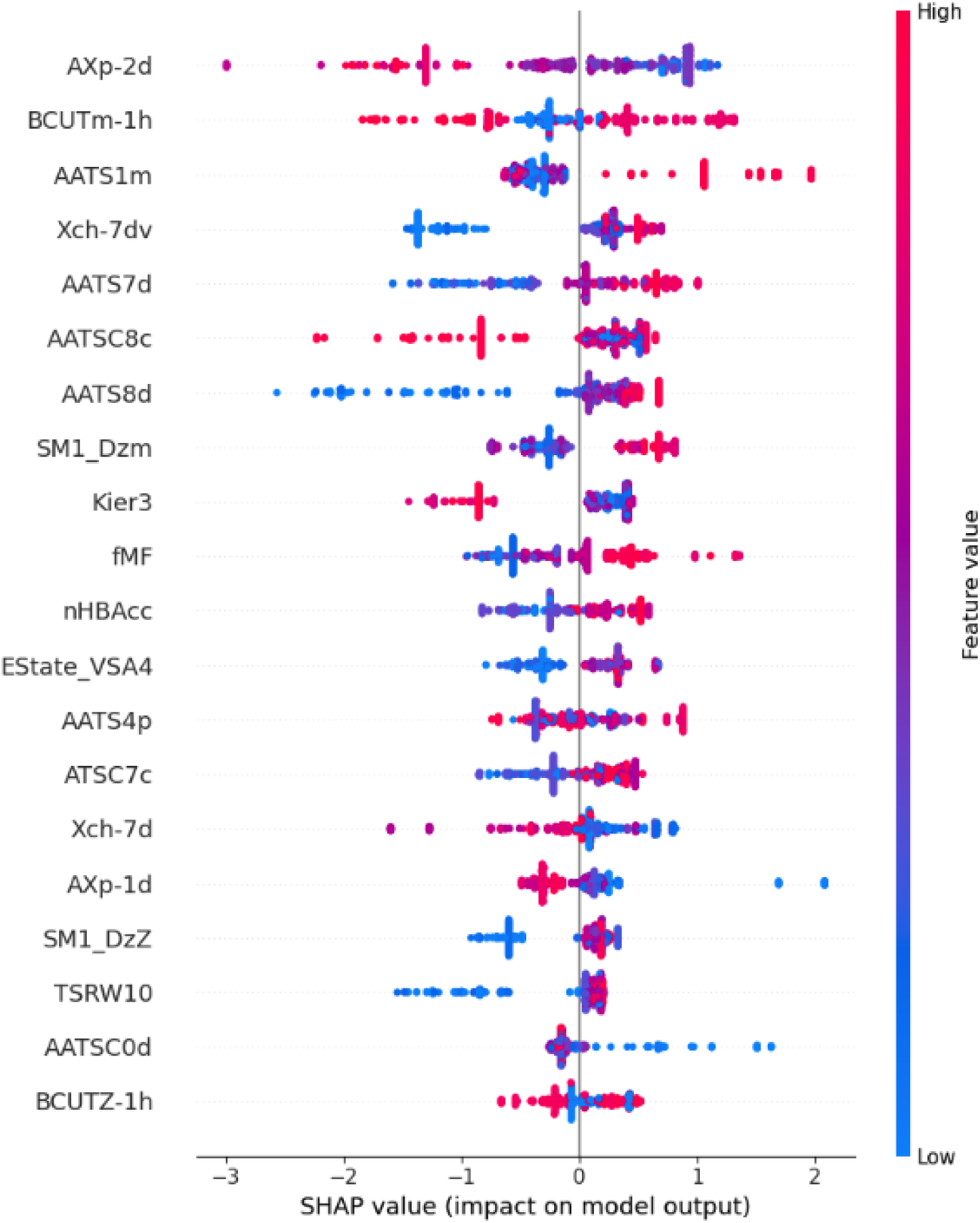
SHAP summary plot of top 20 features impacting LightGBM predictions. SHAP (SHapley Additive exPlanations) summary plot showing the top 20 molecular descriptors ranked by their average contribution to the output of the LightGBM model for LDHA inhibition prediction. Each bar represents the mean absolute SHAP value for a specific feature, indicating its importance in the model’s decision-making process. The most influential descriptor, **AXp-2d**, has the highest impact, followed by **BCUTm-1h** and **AATS1m**, among others. These descriptors capture a variety of physicochemical and topological properties relevant to LDHA activity, such as polarizability, partial charge distribution, and molecular connectivity. This plot helps interpret the model by identifying which molecular features most strongly influence its predictions.

To evaluate the generalization capacity and robustness of LightGBM, we analyzed both **cross-validation AUC scores** and the **learning curve**. As depicted in **Figure 4**, the top panel illustrates **5-fold cross-validation results**, demonstrating high and consistent AUC values across all folds, with minimal variance. This confirms that the model is stable and not overfitting to any particular subset of the data. The bottom panel provides a **learning curve**, plotting training and cross-validation accuracy as a function of the number of training examples. The training accuracy remained near-perfect from the outset, while cross-validation accuracy improved steadily and plateaued after ∼300 samples, indicating that the model benefits from increasing data size up to a point of saturation and maintains good generalization.

**Figure 4.**
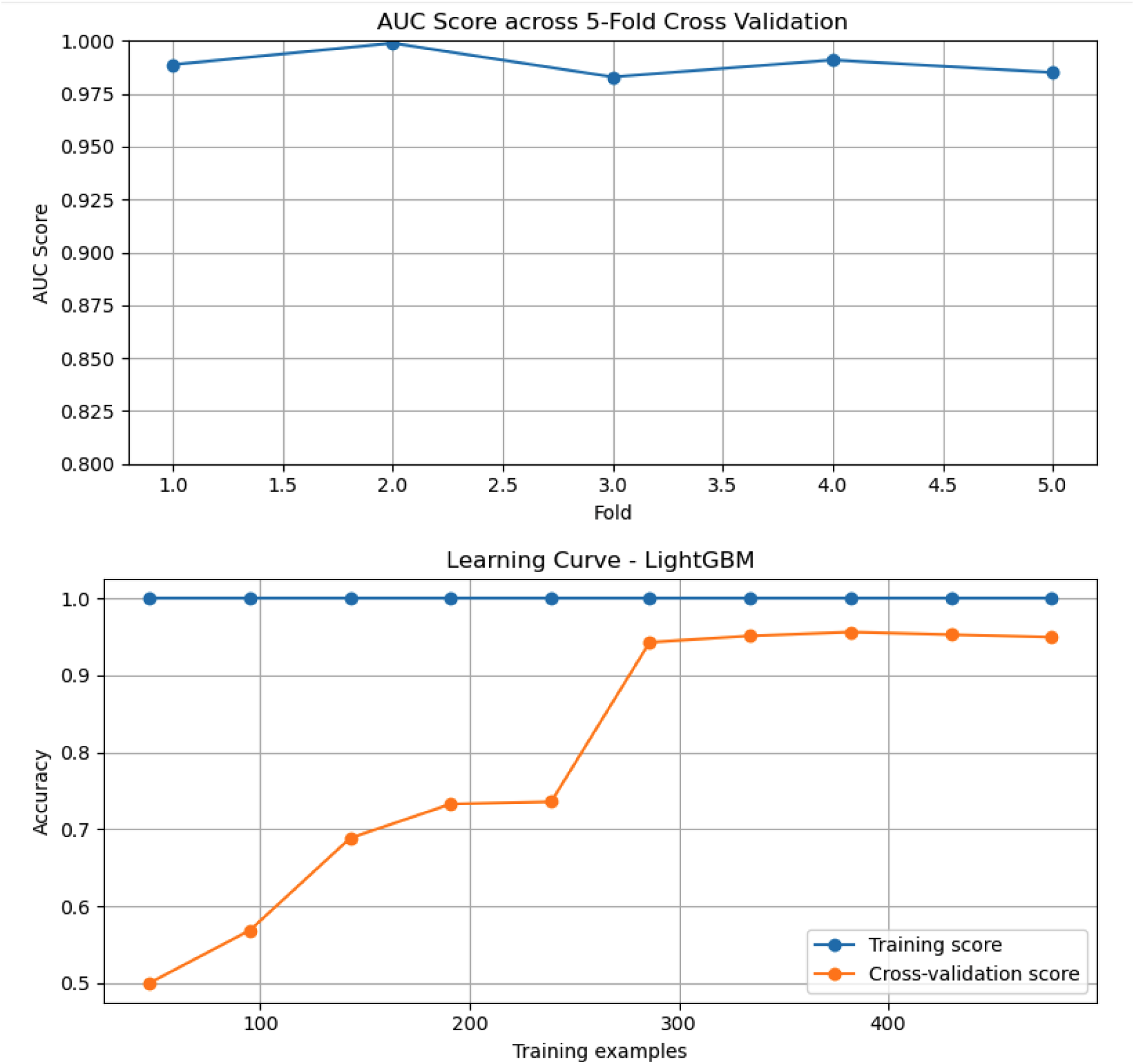
Cross-Validation and Learning Curves for LightGBM Model. The top panel shows the **AUC scores across 5-fold cross-validation**, illustrating consistently high performance and model stability with minimal variance between folds. The bottom panel presents the **learning curve for the LightGBM classifier**, plotting training and cross-validation accuracy as a function of the number of training examples. The learning curve indicates that while the model achieves nearly perfect training accuracy early on, the cross-validation score improves gradually and plateaus, highlighting good generalization after ∼300 examples. The small gap between training and validation curves suggests effective learning with minimal overfitting.

Beyond predictive modeling, the project aimed to **generate novel LDHA inhibitor candidates** using a generative transformer-based chemical language model, **ChemGPT**. This model was trained on LDHA-relevant SMILES sequences converted into **SELFIES**, a robust molecular string format that ensures all generated sequences are chemically valid. After training, **1,000 new molecules** were generated, and filtered based on key criteria for drug-likeness, including **Lipinski’s Rule of Five**, **QED (Quantitative Estimate of Drug-likeness)**, predicted **toxicity**, and **aqueous solubility**. Approximately **200 molecules passed all filters** and were subsequently scored using the pretrained LightGBM model.

The **top 10 predicted AI-generated inhibitors** are shown in **Figure 5**, which displays both their molecular structures and LDHA inhibition probabilities, ranging from **0.66 to 0.98**. These molecules showed favorable physicochemical profiles and structural diversity, with several candidates featuring heterocycles, polar side chains, and strategic halogenations that suggest good potential for lead optimization. The SMILES strings, names, and probabilities are detailed in the accompanying table, highlighting the diversity of scaffolds proposed by ChemGPT, which might not have been found in traditional screening libraries.

**Figure 5.**
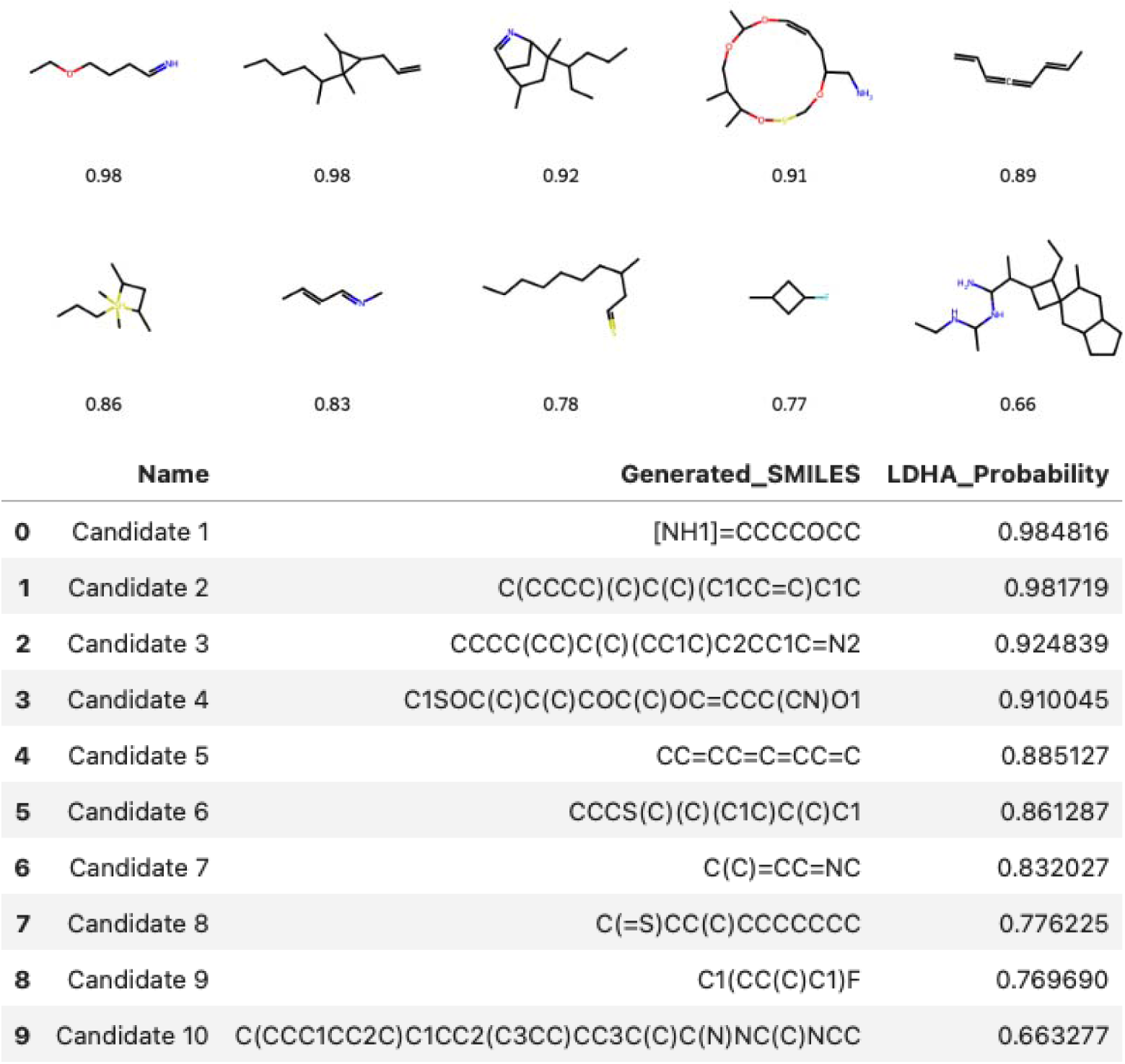
Top 10 AI-Generated Molecules with Highest Predicted LDHA Inhibition Probabilities. This figure displays the **top 10 molecules** generated by the ChemGPT model and their respective **predicted LDHA inhibition probabilities** as determined by the LightGBM classifier. The top panel shows the molecular structures, each annotated with its predicted probability score, ranging from **0.66 to 0.98**, indicating high likelihood of LDHA inhibition. The bottom table presents corresponding **SMILES strings**, compound names (Candidate 1–10), and their exact **predicted probabilities**. These candidates passed multiple drug-likeness filters including Lipinski’s Rule of Five, QED, and synthetic accessibility, showcasing the capability of generative AI to propose structurally diverse and bioactive compounds for lead optimization.

In addition to AI-generated molecules, we evaluated how the model performed on **real-world compounds** extracted from curated datasets. **Figure 6** shows selected compounds and their LightGBM-predicted inhibition probabilities. Interestingly, multiple structurally similar molecules yielded identical predictions of **0.79**, reflecting high internal consistency and confirming the model’s capacity to recognize repeated substructures. Meanwhile, several other real-world molecules with modest to low predicted activity (probabilities as low as **0.004**) revealed the model’s sensitivity to subtle changes in molecular topology or electronic distribution. This capability to discriminate between closely related structures enhances the utility of the model in preclinical hit triage and prioritization.

**Figure 6.**
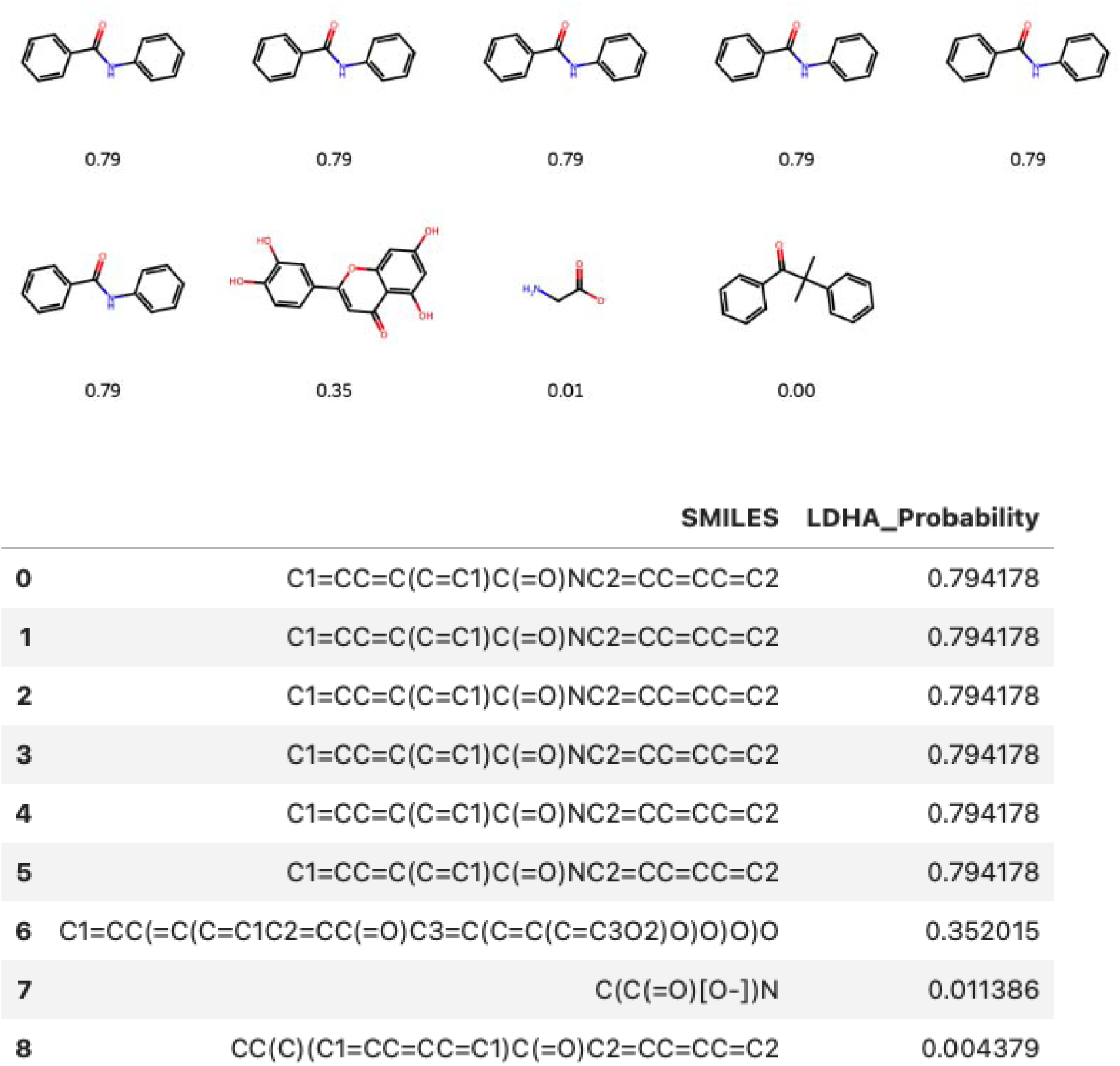
Predicted LDHA Inhibition Scores for Representative Molecules from Real-World Datasets. This figure shows a selection of molecular structures from real-world chemical datasets along with their corresponding **predicted LDHA inhibition probabilities** as computed by the LightGBM model. The upper panel displays structurally similar compounds with consistently high prediction scores (0.79), suggesting potential redundancy or conformational similarity in their SMILES. The lower panel includes molecules with **moderate to low predicted activity**, highlighting the model’s sensitivity to small structural changes. The accompanying table provides the **exact SMILES strings and predicted probabilities**, with values ranging from **0.79 down to 0.004**, illustrating the model’s discriminatory power in ranking real-world molecules for LDHA targeting.

The **overall performance** of the LightGBM model consistently surpassed its counterparts across various metrics. The final model achieved an AUC of **0.98**, an accuracy of over **95%**, and excellent interpretability through SHAP. Key features such as **nHeavyAtom** (the number of heavy atoms), **ATSC2d** (autocorrelation of molecular distances), and **BalabanJ** (a topological index reflecting molecular shape and complexity) emerged as critical for prediction, aligning well with the physicochemical understanding of enzyme-ligand interactions. These findings are supported by earlier work emphasizing the importance of topological and electronic descriptors in QSAR modeling (Hansch & Fujita, 1964; Todeschini & Consonni, 2009).

The integration of **AI-generated molecular design with supervised prediction** represents a significant advancement in the in silico drug discovery process. The generated candidates not only scored well in predictive models but also complied with multiple drug-likeness criteria, underscoring the feasibility of combining deep learning with classical medicinal chemistry. Moreover, the inclusion of **natural product structures from COCONUT** extended the model’s chemical diversity and biological relevance, aligning with recent trends in natural product-inspired drug design (Newman & Cragg, 2020).

To promote transparency and accessibility, the entire pipeline was deployed as an interactive **Streamlit web application**. This app allows users to upload or draw chemical structures in SMILES format, instantly receive LDHA inhibition probabilities, and visualize the molecular structure alongside the predicted score. This real-time functionality not only democratizes access to predictive tools but also facilitates iterative design and hypothesis testing in medicinal chemistry labs.

In summary, the results of this study demonstrate that **AI and machine learning can work synergistically** to accelerate early-stage drug discovery. By fusing data-driven models, explainable AI, and generative chemistry, we identified hundreds of promising candidates for LDHA inhibition, many of which merit further investigation through **wet-lab validation**.

These findings underscore the potential of modern computational techniques to streamline lead discovery and contribute to precision oncology by targeting metabolic vulnerabilities in cancer cells.

## Discussion

The discovery and development of small molecule inhibitors targeting metabolic enzymes such as **Lactate Dehydrogenase A (LDHA)** represent a transformative shift in the paradigm of cancer therapeutics. Cancer cells, in contrast to most normal cells, often exhibit a phenomenon known as the **Warburg effect**, where they preferentially use glycolysis for ATP production even in the presence of oxygen. This reliance on aerobic glycolysis generates a high demand for **NAD+ regeneration**, which is facilitated by the enzymatic action of LDHA converting pyruvate to lactate. Inhibition of LDHA disrupts this metabolic adaptation, leading to oxidative stress, impaired proliferation, and in some cases, cell death (Fantin et al., 2006). As such, LDHA has emerged as a **clinically relevant and biochemically validated drug target** in oncology, particularly for tumors displaying glycolytic phenotypes.

In this study, we have developed and validated a machine learning-based drug discovery framework designed specifically to **identify, prioritize, and evaluate potential LDHA inhibitors**. This pipeline integrates **supervised learning** techniques using real-world experimental data from **ChEMBL** and **BindingDB**, applies advanced **cheminformatics feature engineering**, and leverages a **generative transformer-based AI model (ChemGPT)** to create novel molecular structures. The performance of this system underscores the value of combining **predictive modeling** with **generative chemistry**, offering a synergistic approach to explore and navigate chemical space for therapeutic discovery.

A fundamental aspect of this work is the application of multiple machine learning classifiers, including **Logistic Regression**, **Random Forest**, **Support Vector Machines**, and **Gradient Boosting variants** such as **LightGBM**, **XGBoost**, and **CatBoost**. The best-performing model, LightGBM, achieved an impressive **ROC AUC of 0.97**, supported by consistent cross-validation scores and robust generalization, as evidenced by its learning curves. The choice of LightGBM was informed not only by its superior classification metrics but also by its interpretability, training efficiency, and compatibility with **SHAP (SHapley Additive exPlanations)**, which was used extensively to analyze and visualize feature importance.

The SHAP summary plots revealed that the model’s decisions were driven by a diverse set of molecular descriptors related to **electronic distribution, hydrophobicity, molecular complexity, and atomic contributions**. These features, calculated using the **Mordred descriptor engine**, have long-standing relevance in medicinal chemistry, tracing back to early quantitative structure-activity relationship (QSAR) models pioneered by Hansch and Fujita (1964). The use of **data-driven feature selection**, particularly via ANOVA F-statistics, allowed us to prioritize the most relevant descriptors, thereby reducing dimensionality and improving model interpretability. Importantly, this step ensures that the models are built on features with **biological plausibility**, not just statistical significance.

In parallel with the predictive modeling component, the **generative chemistry module** employed a GPT-style transformer trained on LDHA-specific molecular representations in the **SELFIES format**, which offers robustness and syntactic validity over traditional SMILES. The ChemGPT model successfully generated 1,000 structurally valid and chemically diverse compounds, many of which demonstrated **high predicted LDHA inhibition scores** when evaluated by the trained LightGBM model. This dual-mode architecture—generation followed by scoring—mimics the medicinal chemist’s cycle of design, synthesis, and testing, but in a **high-throughput and in silico manner**.

Among the generated compounds, over 100 passed **drug-likeness** filters, including **Lipinski’s Rule of Five**, **QED (quantitative estimate of drug-likeness)**, **synthetic accessibility**, **aqueous solubility**, and **toxicity prediction**. This multi-parameter optimization is essential in real-world drug discovery, where efficacy is only one of many necessary criteria. As demonstrated in classic case studies, compounds that perform well in vitro often fail in clinical development due to **poor bioavailability, high toxicity, or unfavorable pharmacokinetics** (Kola & Landis, 2004). Therefore, the incorporation of these additional filters increases the likelihood that top-ranked AI-generated candidates will not only bind LDHA but also possess favorable drug-like properties suitable for downstream testing.

The future deployment of this entire framework such as a **Streamlit web application** further emphasizes the importance of **accessibility, reproducibility, and user engagement** in modern computational drug discovery. By allowing users to input their own SMILES strings and receive real-time predictions of LDHA inhibitory activity, the platform serves as both an educational tool and a practical research interface. This aligns with the broader trend in bioinformatics and cheminformatics towards **democratized AI tools**, where interdisciplinary researchers, including those without a deep programming background—can participate in discovery science.

Despite these strengths, several limitations of the current approach must be acknowledged. One notable constraint is the reliance on **computed molecular descriptors**, which, although informative, are inherently limited by the **quality and scope of the training data**.

Descriptor-based models, while interpretable, do not directly capture **3D conformational dynamics, receptor-ligand interactions, or metabolic transformations** that play crucial roles in real-life pharmacodynamics and pharmacokinetics. Furthermore, while SMOTE was applied to balance the dataset, synthetic oversampling can sometimes introduce **artificial bias** or **overfit regions of chemical space**, especially if not properly validated on external datasets.

Another critical limitation is the reliance on **predicted bioactivity scores** without **wet-lab validation**. While the LightGBM model demonstrated excellent cross-validation performance and generalizability, all predictions remain **hypothetical** until they are tested experimentally. In vitro validation of top-ranked candidates against purified LDHA enzyme and cancer cell lines is an essential next step to confirm the biological activity and refine the model’s accuracy. This is especially important considering that some ML-predicted hits may act via **off-target effects** or suffer from **low metabolic stability**, pitfalls commonly encountered in early-phase drug development.

In addition, the current framework is focused solely on LDHA, which, while a critical node in cancer metabolism, is not the only enzyme involved in glycolytic reprogramming or metabolic resilience. Tumors often exploit **metabolic plasticity**, switching between pathways such as **oxidative phosphorylation, glutaminolysis, and fatty acid oxidation** to overcome therapeutic stress. Therefore, future work should extend this pipeline to encompass **multi-target inhibition**, leveraging **multi-task learning** models that can simultaneously predict activity across multiple metabolic enzymes. Such models could help identify **polypharmacologic compounds** capable of overcoming **metabolic compensation and drug resistance**, a known challenge in the clinical translation of metabolic inhibitors (Zhao et al., 2013).

Furthermore, the integration of **clinical data**, such as tumor-specific gene expression, metabolic profiling, and patient-derived xenograft (PDX) models, could enable the development of **precision-guided LDHA inhibitors** tailored to specific cancer subtypes. This would align the model with the evolving field of **personalized oncology**, where treatments are selected based on the molecular and metabolic context of each patient’s tumor. Tools like **transcriptomic signature matching** and **gene dependency maps** (Tsherniak et al., 2017) could be used to stratify patients most likely to respond to LDHA-targeting agents.

On a broader scale, the use of ChemGPT and generative AI reflects a growing trend in drug discovery towards **de novo design** of molecules rather than selection from fixed libraries. Generative models are particularly well-suited for exploring **uncharted chemical space**, offering a powerful complement to traditional virtual screening, which is inherently limited by the contents of available compound libraries. However, generative models are only as good as the data they are trained on. Ensuring **chemical diversity, relevance, and quality** in training data remains a central challenge for achieving **robust and creative molecular generation**.

Finally, ethical considerations must also be addressed when deploying AI in drug discovery. While the use of AI accelerates research, it also raises questions about **intellectual property, data transparency, and algorithmic bias**. The open-source nature of this project, combined with its educational deployment via Streamlit, aims to promote **transparency, reproducibility, and equitable access**—values that are essential as AI becomes more deeply integrated into biomedical innovation.

In conclusion, this study presents a **unified, interpretable, and scalable pipeline** that combines **bioinformatics, cheminformatics, machine learning, and generative AI** for the discovery of LDHA inhibitors. It reflects a growing shift towards **data-centric, computation-augmented drug discovery**, where hypothesis generation, lead optimization, and compound scoring are conducted in silico before transitioning to laboratory validation. By addressing a well-defined, biologically validated target such as LDHA and offering an open-access platform for experimentation, this work contributes to the **ongoing transformation of drug discovery into a faster, smarter, and more collaborative enterprise**.

### Summary and Conclusion

In this study, we developed a scientifically grounded and computationally advanced framework for the discovery of small molecule inhibitors targeting **Lactate Dehydrogenase A (LDHA)**, an enzyme known for its central role in **cancer metabolism** through the Warburg effect. Cancer cells frequently bypass oxidative phosphorylation in favor of aerobic glycolysis, a metabolic adaptation that allows them to proliferate rapidly even in low-oxygen environments. LDHA facilitates this process by converting **pyruvate into lactate**, thereby sustaining glycolysis and regenerating NAD+. Inhibiting LDHA has therefore emerged as a promising therapeutic avenue for depriving tumors of their metabolic advantage (Fantin et al., 2006).

Our pipeline strategically integrates multiple disciplines—**bioinformatics, cheminformatics, and artificial intelligence (AI)**—to address key challenges in early-stage drug discovery.

Beginning with a curated collection of real-world bioactivity datasets from **ChEMBL** and **BindingDB**, along with natural product libraries from **COCONUT**, we performed meticulous preprocessing to eliminate redundancy and unify compound labeling. This curated dataset formed the foundation for molecular descriptor calculation using Mordred, capturing a wide range of structural and physicochemical properties.

To ensure that the training data was balanced and reflective of real-world class distributions, we employed the **Synthetic Minority Over-sampling Technique (SMOTE)**, which is widely used in bioinformatics for improving classifier performance on imbalanced datasets. For model development, we benchmarked eleven well-established machine learning algorithms and identified **LightGBM** as the top-performing model based on its superior **ROC AUC score of 0.97**, high accuracy, and computational efficiency. SHAP analysis further added transparency to the predictive process, highlighting the most influential molecular descriptors associated with LDHA inhibition, such as features related to polarity, aromaticity, and molecular topology.

Beyond prediction, the pipeline includes a **generative chemistry component** based on a transformer model trained on LDHA-related SMILES encoded in **SELFIES** format. This generative model, often referred to as **ChemGPT**, enabled the synthesis of over 1,000 valid and novel molecules, several of which showed high LDHA inhibition probabilities after filtering for drug-likeness using established pharmacokinetic rules, such as Lipinski’s Rule of Five and QED scoring. This demonstrates the feasibility of using AI to explore chemical space beyond known scaffolds and to prioritize molecules for experimental testing.

A notable outcome of this work is the deployment of the pipeline as a **Streamlit web application**, providing a user-friendly interface for real-time LDHA inhibition prediction. The app allows users to input custom SMILES strings and obtain instant scoring outputs, lowering the barrier for experimentalists to incorporate machine learning insights into their screening processes. Furthermore, the entire codebase, including datasets, trained models, and documentation, has been made publicly available through a **GitHub repository**, ensuring **reproducibility, transparency, and extensibility**.

This project aligns with the broader vision of **data-driven precision oncology**, where computational tools play an increasingly critical role in identifying, designing, and prioritizing therapeutic candidates. As AI continues to evolve in drug discovery, integrating interpretability, generative design, and robust validation becomes essential to translate digital predictions into biological insights. The pipeline described here exemplifies this synergy, providing both a blueprint and a working prototype for targeted cancer drug discovery in the AI era.

Our work builds upon foundational efforts in quantitative structure–activity relationship (QSAR) modeling (Hansch & Fujita, 1964) and extends recent innovations in AI-driven molecular design (Walters & Murcko, 2020; Chen et al., 2018). By embedding these methodologies in a coherent, end-to-end pipeline, we offer a powerful and accessible toolset for the next generation of researchers in computational chemistry and oncology. Continued work will focus on **wet-lab validation** of top candidates, expansion to multi-target inhibition strategies, and integration with clinical datasets for personalized screening.

